# The burden of cutaneous and mucocutaneous leishmaniasis in Ecuador (2014-2018), a national registry-based study

**DOI:** 10.1101/751446

**Authors:** Aquiles R. Henríquez-Trujillo, Marco Coral-Almeida, Manuel Calvopiña Hinojosa

**Affiliations:** One Health Research Group, Faculty of Health Sciences, Universidad de Las Américas, Quito, Pichincha, Ecuador

**Keywords:** Cutaneous leishmaniasis, DALY, burden of disease, spatial epidemiology, Ecuador

## Abstract

**Background:** Cutaneous (CL) and mucocutaneous (MCL) leishmaniasis remain as endemic tropical diseases in several Latin American countries. This study aimed to estimate the burden of CL and MCL in Ecuador for the period 2014-2018, in order to inform decision-making and resource allocation to tackle this neglected disease.

**Methods:** Ambulatory consultations, hospitalizations, and reported cases of Leishmaniasis registered by the Ecuadorian National Institute of Statistics and Census and the Ministry of Public Health were used to estimate the burden of CL and MCL during a five-year period. Case estimations were stratified by prevalence of acute and long-term sequelae, to calculate Years Lived with Disability (YLD) by sex and age group using the DALY package in R. Spatial analysis was conducted to identify statistically significant spatial clusters of leishmaniasis.

**Results:** Between years 2014 and 2018, a total 6,937 cases of leishmaniasis were registered, with an average of 1,395 cases reported per year, 97.5% of them were CL and 2.5% MCL. The average cumulative incidence for the study period corrected for underreporting was estimated in 21.98 to 36.10 per 100 thousand inhabitants. Health losses due to leishmaniasis reach 0.32 DALY per 100,000 people per year (95% CI 0.15 – 0.49). The most affected by the disease were men between 15 to 64 years old living below 1,500 m.a.s.l. in sub-tropical and tropical rural communities on both slopes of the Andes mountains. Cantons with the highest cumulative incidence of CL and MCL were Pedro Vicente Maldonado, San Miguel de Los Bancos, and Puerto Quito, in the Pichincha Province; Taisha and Aguarico in the Morona Santiago and Orellana provinces respectively.

**Conclusion:** Compared to previous reports, in the past five years CL and MCL persist as a public health problem in Ecuador. There is a need for more comprehensive and robust data sources to track leishmania cases in Ecuador.

**Author summary:** Cutaneous (CL) and mucocutaneous (MCL) leishmaniasis remains as an endemic neglected tropical disease in several Latin American countries, including Ecuador. Both CL and MCL can produce disfiguring lesions on exposed parts of the body like face and extremities, and permanent scars, contributing to the burden of the disease due to stigma. In order to inform health authorities in their efforts to improve the control of the transmission of the Leishmaniases in the Ecuadorian population, we estimate the burden of CL and MCL for Ecuador in the period 2014-2018, calculating the years lived with disability due to acute and chronic sequelae. We also look for geographical regions within Ecuador with significant clusters of people with the disease, and we found 17 spatial clusters in sub-tropical and tropical rural communities below 1,500 m.a.s.l. on both slopes of the Andes mountains.

## Introduction

The leishmaniases are a group of zoonoses caused by the protozoa of the genus *Leishmania*, which is transmitted by the bite of an infected female sand fly. Around 20 *Leishmania* species are implicated in human disease, producing several clinical presentations from cutaneous (CL) to the destructive mucosal (ML), mucocutaneous (MCL), and a visceral (VL) known as kala-azar, which is often fatal if left untreated [1, 2].

The republic of Ecuador is a country located in South America, crossed transversally by the Equator line and longitudinally (north to south) by the Andes belt mountains, dividing the country in four natural regions. Each region has different ecosystems, but also attitudes, believes and life style of the people. These ecosystems include richest in flora and fauna for transmission of vector-borne parasitic diseases including *Leishmania* at each region. *Leishmania* vectors and reservoirs showed a broad distribution across all the biomes of Ecuador, with higher risk in the lowlands [3].

In Ecuador CL and MCL are considered a public health problem by the Ministry of Public Health (MPH), and the disease is of the mandatory notification in the country since 2005. Leishmaniases affect mainly children in rural areas of the 3 natural regions, with a wide range geographic distribution, affecting 23 of the 24 provinces, except Galapagos Islands [4]. Regarding occurrence of clinical forms, CL is the most common followed by MCL. CL is considered self-healing in a period of 9 months but MCL never cure itself, with chronic progression [2]. Clinical variants of CL are recidiva cutis, diffuse-CL, disseminated-CL, nodular, verrucoide, sporotricoid, amongst others [4]. Until now, not a single case of VL has been confirmed in Ecuador.

In the country, both *Leishmania* and *Viannia* subgenus has been identified with a total of 8 species. *L. (V.) braziliensis* predominate in the Amazon region, *L. panamensis/guyanensis* in the subtropical and tropical lowlands of Pacific region, whilst *L. (L.) mexicana* in the inter-Andean valleys [5]. Most of the MCL cases are infected in the Amazon region associated with the most virulent *L. braziliensis* [6]. Regarding the proportion of cases of CL and MCL, only 6.9% (18/260 cases) showed the destructive form [7]. However, in an active survey13 cases of MCL were diagnosed in the Amazon region [8]. In another study reporting from different regions, a total of 148 Leishmania-positive subjects, revealed that 135 (91.2%) were CL and the rest 13 (8.8%) MCL [4]. In subtropical Pacific side of Pichincha province, 432 clinical records of leishmaniasis-patients recorded during five years (2007–2011) were analyzed, all represented CL lesions [9].

The clinical features of CL are mainly ulcers, followed by chiclero’s ulcers, recidiva cutis and nodular lesions [5]. The MCL cases presented with erythema, ulcerations, granulomas, septal perforation, swelling of upper lip and nose, bleeding and crusts [5]. CL is considered a self-cure disease but not MCL. However, the clinical form recidiva cutis could still active and growing for several years [10]. CL lesions leaves a permanent atrophic scar, some studies confirmed high prevalence of scars in endemic areas [11]. In a study in the subtropical area 75% of patients with CL cured without treatment after 9 months of follow-up [12].

Disability-adjusted life years (DALY) is a synthetic measure of disease burden that has been widely used to measure the global burden of disease since 1991[13]. DALY combines the years of life lost due to premature death (YLL) and the years lived with disability (YLD) caused by the disease and can be interpreted as healthy life years lost [14, 15]. Globally, Andean Latin American countries are among those with a high proportion of healthy life years lost due to CL and MCL [16]. This study aimed to estimate the burden of cutaneous and mucocutaneous leishmaniasis in Ecuador during the period 2014-2018, in order to inform decision-making and resource allocation to tackle this neglected tropical disease.

## Methods

### Geographical location and study population

Ecuador is located in the Pacific coast of South America, limiting to the North with Colombia and to the South and East with Perú. The country has 283,561 km² and four geographical regions: The Galapagos Islands, the coastal region, the highlands of the Andean mountains, and the Amazonia. The climate of each region is largely determined by altitude, with humid subtropical and tropical weather on western and eastern slopes of the Andes below 1500 m.a.s.l., and dry temperate weather in the inter-Andean valleys above 1500 m.a.s.l. For administrative purposes the country is divided into 24 provinces, and 224 cantons. The last nation-wide census in year 2010 recorded 15 million inhabitants, and the projected population for year 2019 is 17.2 million inhabitants [17].

### Sources of information and study design

This is a national cross-sectional registry-based study, using all the information available from the registries of ambulatory consultations, hospitalizations, and reported cases of leishmaniasis registered by the Ecuadorian National Institute of Statistics and Census (INEC) [18], and the Ecuadorian Ministry of Public Health (MPH) [19].

### Burden estimations

To estimate the burden of disease of CL and MCL during a five-year period, cases were identified by their International Statistical Classification of Diseases 10th revision (ICD-10) codes: B55.1 for Cutaneous leishmaniasis, B55.2 for Mucocutaneous leishmaniasis, and B55.9 for unspecified Leishmaniasis [20]. Case frequencies were adjusted for underreporting using expansion factors between 2.8 to 4.6 fold based on previous estimations published by Alvar J. et al. [21]. Case estimations were stratified by prevalence of acute and long-term sequelae, to calculate estimated cumulative incidences. Burden was measured in Disability adjusted Life Years (DALY), which are calculated by the sum of Years Lived with Disability (YLD) and years of life lost due to premature mortality (YLL) [14]. We assumed that mortality due to CL and MCL was null, therefore DALY values are the same as for YLD following the methods described by Karimkhani C. et al. [16]. Disability weights (DW) used for short and long term sequalae were obtained from the estimates published by the Global Burden of Disease 2013 study [22].The DW used for the short term sequalae of cutaneous leishmaniasis were those related to “Disfigurement: level 1 = 0.011 (95%CI 0.005 to 0.021)”, and for long term sequalae mainly related to mucocutaneous leishmaniasis “Disfigurement: level 2 = 0.067 (95%CI 0.044 to 0.096)”. YLD by sex and age group were calculated using the DALY package in R considering a time discount rate of 3% per year, without age weighting [14]. To estimate the duration of long term sequalae we considered the residual life expectancy from the Coale and Demeny model life table West 26 with a life expectancy at birth of 80 years for males and 82.5 years for females [23].

### Spatial analysis

Spatial analysis was conducted in SaTScan™ version 9.6 to identify statistically significant spatial clusters of Cl and MCL [24]. The spatial unit for the analysis were cantons, the second-level territorial subdivisions below the provinces. The spatial clusters were defined by the cantons with higher prevalence of ambulatory consultations for cutaneous or mucocutaneous leishmaniasis during the study period. To perform the analysis, the following variables were used: 1) the number of CL+MCL cases distributed geographically by canton, 2) the total population of each canton, and 3) the spatial coordinates of each canton. The count of cases in each geographical location were compared using a Poisson distribution. Space clustering was determined by the contrast of the incidence rate ratio of leishmaniosis cases within an area with an expected incidence rate ratio of the leishmaniosis cases if their incidences were randomly distributed. The SaTScan™ software tested if any clusters with an increased risk for the occurrence for leishmaniasis can be detected using the method described by Kulldorff M. [25]. Likelihood ratio test was used to check the significance of identified space clusters, Monte Carlo simulations with 999 iterations were used to assess the significance of the results of the test (p-values). A cluster was considered as significant when p-values were inferior to 0.05 [26]. The Gini coefficient was used as an additional selection filter amongst the significant clusters following the methodology described by Han et al. [27]. Visualization of the spatial analysis was done using QGIS version 3.8 software [28]. Shape files for the maps in this article were obtained from the INEC portal following their licensing requirements [29]. All maps were created and designed by the authors of this manuscript.

### Ethics statement

Ethical approval was not required for this study. All data used in this study is freely available from public databases of the Ecuadorian National Institute of Statistics and Census (INEC) and the Ecuadorian Ministry of Public Health (MoH) [18, 19]. By legal mandate, all the records from these databases are anonymized, and can be used for research and educational purposes while keeping confidentiality [30].

## Results

Between years 2001 and 2018 the MPH registered 27,095 leishmaniases cases (see Fig 1). In the study period between years 2014 and 2018, there were 6,937 cases notified, with an average of 1,395 cases reported per year, 97.5% of them were cutaneous leishmaniasis and 2.5% mucocutaneous leishmaniasis. The crude cumulative incidence was 8.4 per 100 thousand inhabitants per year (95% credibility interval 6.6 to 10.3). When corrected for underreporting, we estimate that the cumulative incidence for the study period may be in the range between 29.53 to 48.52 per 100 thousand inhabitants (Table 1). On average 60.2% of patients were men. In respect of age, 30.3% of the cases occurred in patients less than 15 years old, 63.7% of the cases in people between 15 to 64 years old, and 6% in persons 65 years old and beyond. In average there were 3 ambulatory consultations per patient. Average health losses due to leishmaniasis reach 0.32 DALY per 100,000 people per year (95% CI 0.15 – 0.49). All the burden was due to Years Lived with Disability.

**Fig 1.**
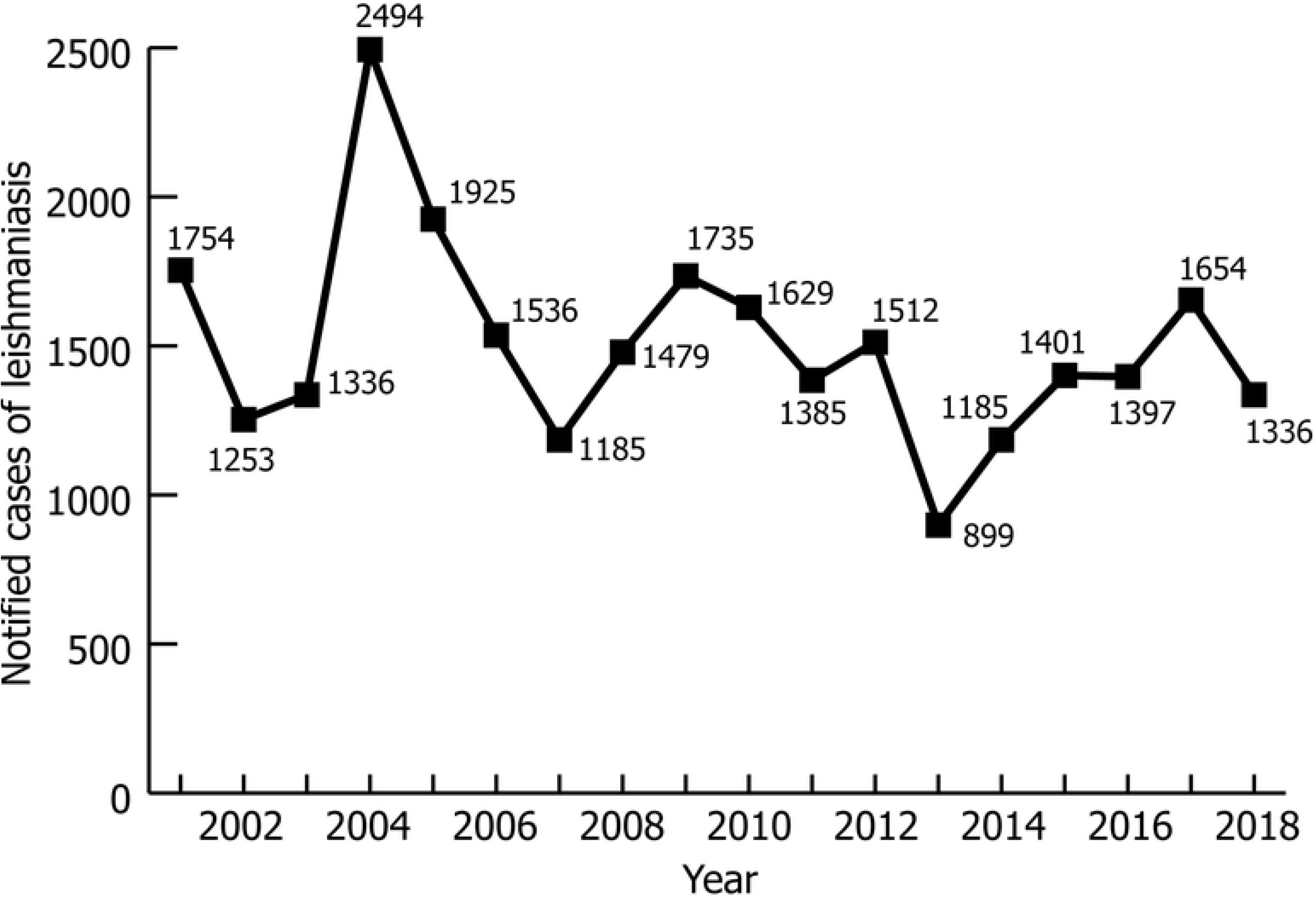
Notified cases of leishmaniases in Ecuador, 2001-2018. Cases of leishmaniases registered in EPI-2 and SIVE-Alerta registries published by the Ecuadorian Ministry of Public Health. Available from: https://www.salud.gob.ec/gaceta-epidemiologica-ecuador-sive-alerta/ [19].

**Table 1.** Cumulative incidence of cutaneous leishmaniasis in Ecuador between years 2014-2018.

| Year | Total cases | CL | MCL | Estimated CL+MCL cases <sup>a</sup> | Incidence rate <sup>b</sup> |
| --- | --- | --- | --- | --- | --- |
| **2014** | 1 185 | 1 185 | NR | 3 318 to 5 451 | 20.70 to 34.01 |
| **2015** | 1 401 | 1 382 | 19 | 3 923 to 6 445 | 24.10 to 39.59 |
| **2016** | 1 397 | 1 397 | NR | 3 912 to 6 426 | 23.67 tot 38.88 |
| **2017** | 1 654 | 1 654 | NR | 4 631 to 7 608 | 27.60 to 45.35 |
| **2018** | 1 336 | 1 302 | 34 | 3 741 to 6 146 | 21.98 to 36.10 |
| **Total** | 6 973 | 6 920 | 53 | 19 524 to 32 076 | 29.53 to 48.52 |

CL = Cutaneous leishmaniasis; MCL = Mucocutaneous leishmaniasis; NR = Not reported
<sup>a</sup> Correction for underreporting (2.8 to 4.6-fold) based on Alvar J. et al.[21]; <sup>b</sup> Incidence rates per 100 thousand inhabitants.

**Table 2.** Estimated DALY due to cutaneous leishmaniasis in Ecuador between years 2014-2018.

| Year | Estimated cases | DALY | DALY rate <sup>a</sup> |
| --- | --- | --- | --- |
| 2014 | 4 385 | 45 (22 to 69) | 0.28 (0.14 to 0.43) |
| 2015 | 5 183 | 53 (25 to 81) | 0.33 (0.15 to 0.50) |
| 2016 | 5 169 | 53 (25 to 81) | 0.32 (0.15 to 0.49) |
| 2017 | 6 155 | 63 (30 to 96) | 0.38 (0.18 to 0.57) |
| 2018 | 4 943 | 51 (24 to 77) | 0.30 (0.14 to 0.45) |

DALY = Disability-Adjusted Life Years; <sup>a</sup> Rate per 100 thousand inhabitants

As observed in **Fig 2**, the cantons with the with highest number of cases of CL and MCL per 10,000 inhabitants (in the range between 212 to 464 cases per 10,000 inhabitants) were Pedro Vicente Maldonado, San Miguel de Los Bancos, and Puerto Quito, in the Pichincha province on the western slope of the Andes below 1,500 m.a.s.l. In the Amazon region Taisha and Aguarico cantons at the Morona Santiago and Orellana provinces respectively, also below 1,500 m.a.s.l, have a higher number of cases. A total of seventeen significant spatial clusters were distributed in the tropical and subtropical zones of Ecuador (p<0.001). All the cantons in the Amazon region were part of at least one significant cluster for leishmaniasis, 4 clusters were identified in the borderline between the northern and central-northern coastal region of Ecuador and the highlands, and a single cluster was located in the coastal northern province of Esmeraldas. Relative risks of the clusters varied from 1.79 to 48.58.

**Fig 2.**
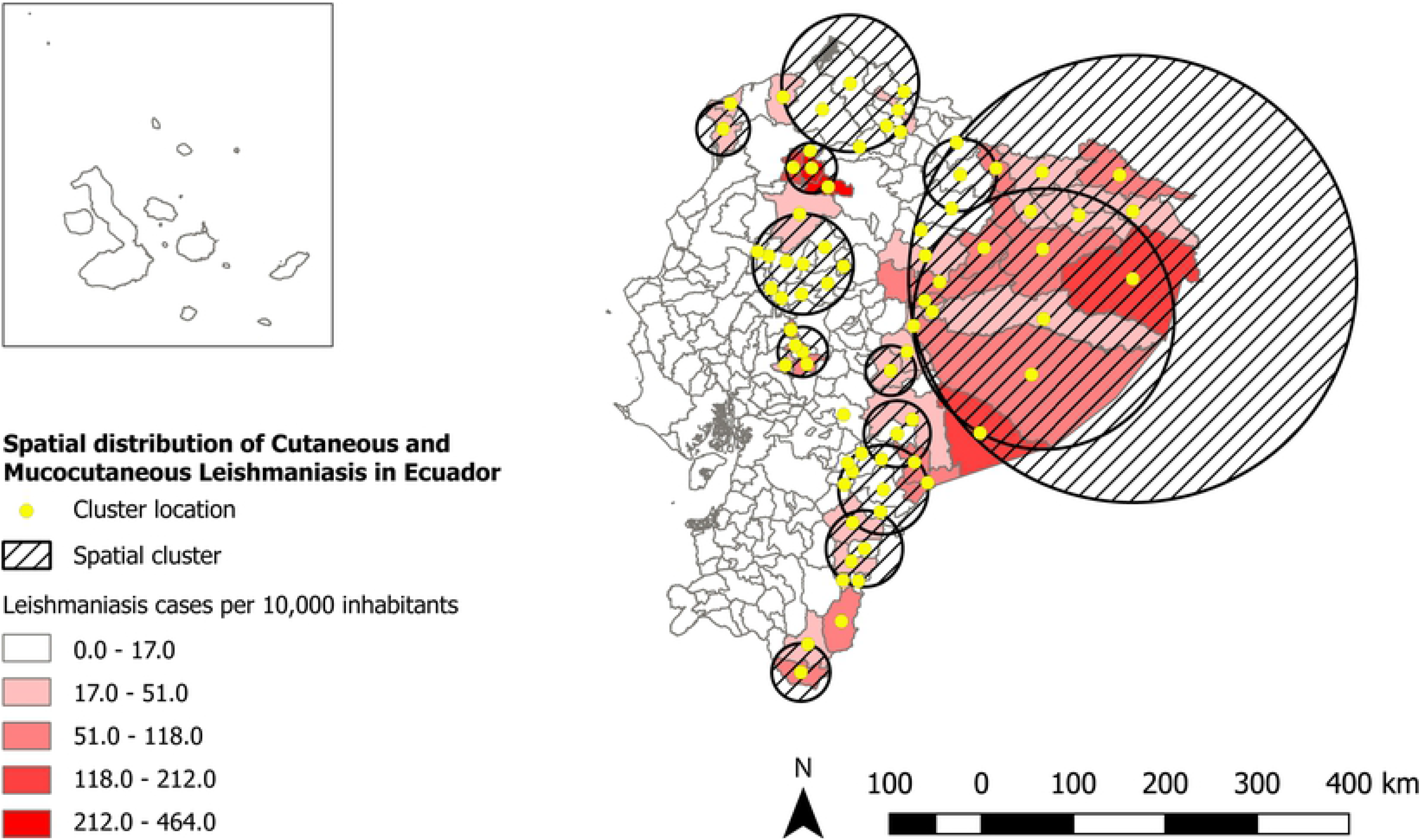
Spatial clusters of cutaneous an mucocutaneous leishmaniasis in cantons from Ecuador, 2014-2018. Spatial clusters of cutaneous and mucocutaneous leishmaniases registered by canton of residence of the patient. Shapefile used for the cantonal limits in the map are freely available after registration in: https://www.ecuadorencifras.gob.ec/registro-de-descargas-cartograficas/ [29].

## Discussion

This is the first report using DALY to estimate the local burden of disease due to cutaneous and mucocutaneous leishmaniasis in Ecuador, and the first to analyze for possible spatial clusters at country level. Furthermore, the present study updates previous epidemiological reports about the epidemiology of leishmaniases in Ecuador [5, 11, 31]. Compared to these local reports, in the past five years CL and MCL persist as a public health problem in Ecuador, and both clinical forms remain endemic. As reported in several studies [16, 21, 31], Andean Latin American countries are among those with a high proportion of healthy life years lost due to CL and MCL. However, the overall frequency of CL and MCL in Ecuador is much lower than reported in Peru and Bolivia.

Our estimates are consistent with the burden reported in global studies [16], but our numbers are closer to the lower limits of published estimates despite correction for underreporting. The cross-sectional analysis of data from the GBD 2013 study published by Karimkhani C. et al. calculated a mean age-standardized burden of CL of 0.58 DALY per 100,000 people worldwide, and estimated 0.96 (0.44 – 1.88) DALY per 100,000 people for Ecuador, with an increase of 2.96% between from year 1990 to 2013 [16]. In our estimations the burden of CL and MCL was lower 0.32 (0.15 – 0.49) DALY per 100,000 people per year. We think this difference is partially due to the influence of the scarcity of data sources and lack of robust registries, generating more uncertainty both in the global estimates and in our estimates. This problem with the quality of the registries was already pointed out by Reithinger R. in a letter published in year 2016 [32].

There is a need for more comprehensive and robust data sources to track leishmania cases in Ecuador. The MPH begun passive surveillance of CL and MCL trough case notification since year 2005 onwards. Despite case notification is compulsory and is made through computerized online systems (SIVE-ALERTA, RDACAA, PRAS, and others), available data comes only from the consultations registered by the public healthcare services; therefore, we considered it represents a subset of the real incidence and prevalence of leishmaniasis at the national level. We must remember that CL in Ecuador is a self-resolving disease in 75% of the cases [12], and many patients may not seek attention. A publication by Alvar J. et al, estimate underreporting of CL being in the range of 2.8 to 4.6 times for the Latin American countries [21]. Without active surveillance, empirical data collection and field validation, any registry will be inaccurate.

Finally, there is a need of a more comprehensive approach, to better understand the complex relationships between individual, social, animal and ecological factors influencing CL and MCL epidemiology, and better inform decision makers for the control of leishmaniasis. Climate variability influences vector distribution, and vector-borne disease transmission. Models published by Escobar L. et al. have predicted a rise in leishmaniasis exposure risk in the Andes Mountains by 2030 and 2050, mainly related to climate warming [33]. Our spatial analysis showed that CL and MCL cases in Ecuador are evenly distributed on both sides of the Andes slopes below 1,500 m.a.s.l. which are regions with sub-tropical and tropical climate. The cantons of the North-West of the Pichincha Province, and those in the Amazon region had a higher risk of CL and MCL when compared to the other regions of the country. It is possible that new important ecological niches are found in this region and should be further explored. The Amazon region is vast and has a low population density when compared to the rest of the country, for these reasons, local surveillance should be applied for leishmaniasis cases and more studies are required regarding the transmission of leishmaniasis. The coastal clusters of leishmaniasis show that there is an uneven distribution in these lowlands, it is necessary to study the cause of these differences in the search of a possible explanation for protective/risk factors associated with leishmaniasis cases.

## Supporting information

**S1 Checklist**. STROBE Checklist for the article.

